# MicroRNA-122-mediated liver detargeting enhances the tissue specificity of cardiac genome editing

**DOI:** 10.1101/2023.06.29.546982

**Authors:** Luzi Yang, Congting Guo, Zhanzhao Liu, Zhan Chen, Yueshen Sun, Xiaomin Hu, Yanjiang Zheng, Yifei Li, Fei Gao, Pingzhu Zhou, William T. Pu, Yuxuan Guo

**Affiliations:** Peking University Health Science Center, School of Basic Medical Sciences, Beijing, 100191, China; Peking University Institute of Cardiovascular Sciences, Beijing, 100191, China; Peking Union Medical College Hospital, Department of Cardiology, Chinese Academy of Medical Science & Peking Union Medical College, Beijing, 100730, China; State Key Laboratory of Complex Severe and Rare Diseases, Peking Union Medical College Hospital, Department of Medical Research Center, Chinese Academy of Medical Science & Peking Union Medical College, Beijing, 100730, China; Ministry of Education Key Laboratory of Birth Defects and Related Diseases of Women and Children, Department of Pediatrics, West China Second University Hospital, Sichuan University, Chengdu, 610041, China; Beijing Anzhen Hospital, Department of Cardiology, Capital Medical University, Beijing, 100029, China; Boston Children’s Hospital, Department of Cardiology, Boston, MA 02115, USA; Harvard Stem Cell Institute, 7 Divinity Avenue, Cambridge, MA 02138, USA; State Key Laboratory of Vascular Homeostasis and Remodeling, Peking University, Beijing, 100191, China; Beijing Key Laboratory of Cardiovascular Receptors Research, Beijing, 100191, China

**Keywords:** adeno-associated virus, cardiac gene editing, microRNA-122, liver detargeting

## Abstract

**Background:** The cardiac troponin T (Tnnt2) promoter is broadly utilized for cardiac specific gene expression, particularly via adeno-associated virus (AAV)-based gene transfer. However, these vectors drive lower-level ectopic gene expression in other tissues, most notably in the liver. Whether the AAV-*Tnnt2* vectors remain tissue-specific in applications sensitive to low or transient gene expression, such as gene editing, remains unclear.

**Methods:** The tissue specificity of AAV9-*Tnnt2* vectors was evaluated in mice using Cre-LoxP-based fluorescence reporters and CRISPR/Cas9-mediated somatic mutagenesis. CRISPR/Cas9-triggered AAV integration into host genome was further assessed by quantitative PCR.

**Results:** In mice treated with AAV-*Tnnt2*-GFP, GFP signal was specifically observed in the heart by confocal imaging. However, when AAV-*Tnnt2*-Cre was administered to mice carrying LoxP-STOP-LoxP fluorescence reporters, the reporter signals were observed in up to 50% hepatic cells. Similarly, the AAV-*Tnnt2*-SaCas9 vector extensively edited the hepatic genome as measured by targeted amplicon-sequencing. Cas9-triggered AAV integration into the host genome was also validated in the liver. Inclusion of target sequences for microRNA-122, a highly expressed, liver-specific microRNA, in the AAV transgene’s 3’ untranslated region (3’ UTR) markedly reduced ectopic transgene expression, genome editing and AAV integration in the liver.

**Conclusions:** The heavily used AAV-*Tnnt2* system exhibits liver leakiness that severely impairs the cardiac specificity of AAV-based genetic manipulation. This problem can be mitigated via miR122-mediated liver detargeting.

---

Recombinant adeno-associated virus (rAAV) is broadly applied in cardiovascular research and gene therapy. While the most widely used rAAV, serotype 9 (AAV9), robustly transduces the liver, the heart, and other organs, its gene expression can be selectively targeted to cardiomyocytes in the heart using a cardiac specific promoter, most commonly that from cardiac troponin T gene (Tnnt2 or cTnT) ^1^. This AAV9-Tnnt2 system is increasingly favored both in the investigation of cardiac disease mechanisms and gene therapy strategies. However, its specificity for the heart relative to other major AAV-targeted organs, especially the liver, requires greater investigation. This issue is especially important for AAV-delivered recombinase-based genome manipulation or CRISPR-mediated genome editing, where low or transient gene expression is sufficient to cause profound biological outcomes.

To assess AAV9-Tnnt2-driven gene expression, we constructed AAV9 vectors AAV-Tnnt2-GFP and AAV-Tnnt2-Cre (Figure A), which share the same vector backbone and only differ in the coding sequences. We injected 5×10^10^ vg/g (vector genome per gram body weight) rAAVs subcutaneously into postnatal day 1 (P1) R26^fsCas9-GFP^ mice (Figure A), which harbor a Cre-activatable GFP reporter. At P14, we evaluated rAAV-mediated transgene expression by GFP fluorescence imaging. In AAV-Tnnt2-GFP treated mice, GFP signal was only detectable in the heart by confocal microscopy (Figure B-C). Strikingly, in AAV-Tnnt2-Cre treated animals, we observed GFP signal in up to 50% cells in the liver (Figure B-C). GFP signal was not detected in spleen, lung, kidney, brain, muscle, or gonads (Figure B-C). AAV-Tnnt2-Cre-triggered LoxP recombination was also confirmed in the liver of R26^fsCas9tdTomato^ mice (Figure B-C), a reporter mouse that was independently generated using gene targeting vectors different from R26^fsCas9-GFP^. The above observation indicated that the commonly used AAV-Tnnt2-GFP assay does not have sufficient sensitivity to report transgene expression in the liver, which can be detected by the Cre-LoxP reporters.

**Figure.**
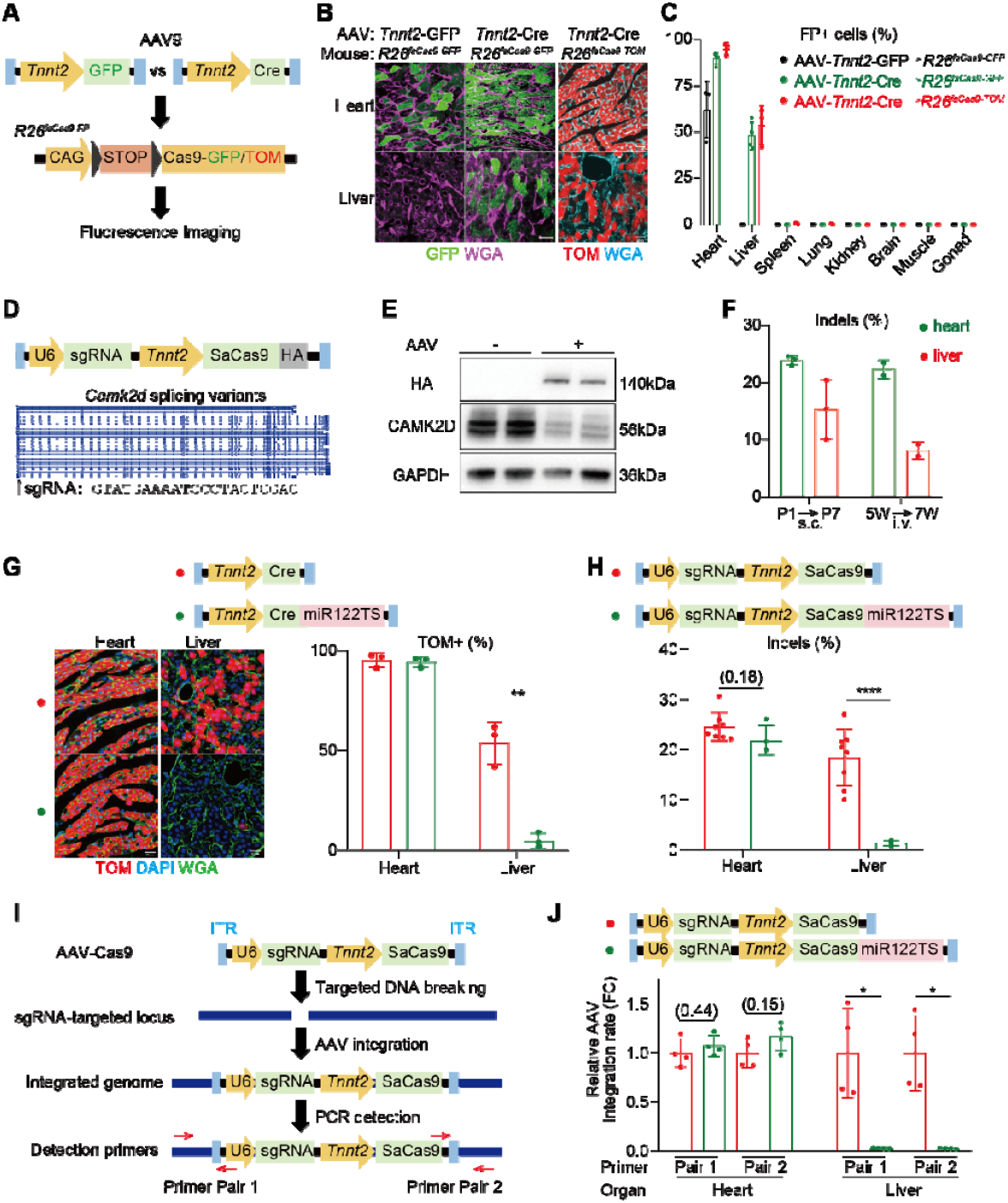
MicroRNA-122 targeting sequence reduces the liver leakage of AAV-*Tnnt2*-based cardiac gene delivery. **A**, the study design to compare AAV transgenic GFP reporter versus AAV-Cre-LoxP fluorescence protein (FP) reporters in the assessment of AAV tissue specificity. TOM, tdTomato. Dark grey triangles, LoxP. **B**, confocal images of tissue cryosections. **C**, quantification of FP-positive cells in the various tissues. **D**, the vector design of AAV-Tnnt2-SaCas9 targeting *Camk2d* exon 2. HA, hemagglutinin tag. **E**, western blot analysis of CAMK2D in AAV-SaCas9-treated hearts. **F**, indel quantification via amplicon-sequencing and CRISPResso2 analysis. AAV injection and sample collection ages labeled below the plots. s.c., subcutaneous; i.v., intravenous. **G**, AAV-Tnnt2-Cre-miR122TS vector design and its impact on Cre-LoxP activation. **H**, AAV-Tnnt2-SaCas9-miR122TS vector design and its impact on gene editing. **I**, a schematic of AAV-Cas9-triggered AAV integration into sgRNA-targeted loci and the primer designs for quantitative PCR detection. ITR, inverted terminal repeats. **J**, qPCR-based analysis of AAV integration into the sgRNA-targeted genome. FC, quantitative PCR fold change. In all panels, scale bars stand for 20 μm. Mean± SD. Student’s t test: *P<0.05, **P<0.01, ****P<0.0001, non-significant P values in parenthesis.

Cre-LoxP is a permanent DNA recording system responsive to low or transient Cre expression. CRISPR/Cas9 gene editing is known exhibit a similar behavior. Thus, we next constructed an AAV-Tnnt2-SaCas9-U6-sgRNA vector to test if AAV-Tnnt2 vectors could cause ectopic gene editing in the liver (Figure D). The backbone of this vector was distinct from the AAV-Tnnt2-GFP/Cre vectors to reduce the likelihood that backbone sequences contribute to the leaky rAAV transgene expression. SaCas9 (*Staphylococcus aureus* Cas9) is a miniature Cas9 homolog that allows all CRISPR/Cas9 components to be delivered by a single AAV vector. We designed a SaCas9 single-guide RNA (sgRNA) targeting the exon 2 of *Camk2d*, an exon shared by all splicing variants of this gene (Figure D). *Camk2d* encodes the major cardiac isoform of calcium/calmodulin-dependent protein kinase II (CaMKII), a heavily studied therapeutic target that requires stringent cardiac specificity for the safe treatment of heart diseases ^2^.

We first subcutaneously applied the AAV-Tnnt2-SaCas9-*Camk2d*-sgRNA vector to P1 wildtype mice and analyzed tissues at P7. Western blot analysis validated the efficient depletion of the CAMK2D protein in the heart (Figure E). Next-generation sequencing of the sgRNA-targeted genomic loci revealed more than 10% DNA insertions and deletions (indels) in the liver, confirming leaky gene editing by the AAV-Tnnt2 system (Figure F). We observed similar results when the vector was intravenously injected into 5-week-old animals (Figure F), thus the AAV-Tnnt2-mediated ectopic gene editing in the liver was independent of the animal ages or the routes of systemic administration.

MicroRNA-122 (miR122) is the most abundant microRNA that is specifically expressed in the liver. Incorporation of the miR122 target sequences (miR122TS) into the 3’ untranslated region (3’ UTR) of AAV transgenes suppressed their expression in the liver ^3, 4^. Thus, we tested if the inclusion of miR122TS could reduce the liver leakage of the AAV-Tnnt2 system. We treated R26^Cas9tdTomato^ mice with AAV9-Tnnt2-Cre-miR122TS and strikingly observed more than 90% reduction of the ectopic tdTomato-positive cells in the liver as compared to mice treated with AAV9-Tnnt2-Cre (Figure G). Similarly, miR122TS also drastically reduced the hepatic gene editing by the AAV-Tnnt2-SaCas9-*Camk2d*-sgRNA vector without altering the cardiac gene editing rate (Figure H). The aforementioned AAV-Tnnt2-Cre-miR122TS and AAV-Tnnt2-SaCas9-miR122TS vectors harbor distinct 3’UTR-PolyA sequences, confirming that the miR122TS function is not dependent on its flanking sequences.

One major safety concern in AAV gene therapy is relevant to vector integration into liver genome. In particular, CRISPR-triggered DNA breaks enhance AAV integration into the sgRNA-targeted loci ^5^ (Figure I). Thus, we next tested if miR122TS could reduce Cas9-mediated AAV genome integration in the liver. We designed PCR assays to specifically detect the boundaries between AAV and host genome DNA (Figure I) and quantitatively assessed AAV integration by qPCR. Strikingly, we observed that the inclusion of miR122TS eliminated AAV integration into liver genome, while cardiac AAV integration was not affected (Figure J). Therefore, miR122TS can avoid the AAV genome integration that is secondary to the AAV-delivered genome editing in the liver.

In summary, this study evaluated the cardiac specificity of the broadly used AAV-Tnnt2 gene delivery system and uncovered extensive transgene leakage in the liver. This technical problem is particularly relevant to AAV-based gene editing, as AAV-Tnnt2-Cas9 vectors not only trigger hepatic mutagenesis but also enhance AAV integration into the liver genome. Fortunately, this problem can be solved by adding miR122TS to the 3’UTR of rAAV transgenes. Thus, the new AAV-Tnnt2-miR122TS system provides a powerful and essential tool to enhance the cardiac specificity of AAV-based cardiovascular research and gene therapy.

## Acknowledgements

We thank PackGene Biotech for AAV production and Novogene for next-generation sequencing.

## Author Contribution

Y.G. conceived the research and supervised the study. L.Y., and P.Z., independently observed the liver leakiness of AAV-Tnnt2 system in two different labs. L.Y., C.G., Z.L. and Z.C. conducted the research and analysis. Y.S. and X.H. assisted in mouse experiments. Y.Z. and Y.L. independently validated the *Camk2d* gene editing results in a different lab. W.T.P. provided advice in AAV genome integration experiments and manuscript revision. Y.G. wrote and revised the manuscript.

